# Rational domestication of a plant-based recombinant expression system expands its biosynthetic range

**DOI:** 10.1101/2021.12.09.472022

**Authors:** Mark A. Jackson, Lai Yue Chan, Maxim D. Harding, David J. Craik, Edward K. Gilding

**Affiliations:** Institute for Molecular Bioscience, Australian Research Council Centre of Excellence for Innovations in Peptide and Protein Science, The University of Queensland, Brisbane, Queensland 4072, Australia

**Keywords:** Plant molecular farming, gene editing, CRISPR-Cas9, asparaginyl endopeptidase, recombinant, protease, cyclotide, peptide, therapeutic, insecticide

## Abstract

Plant molecular farming aims to provide a green, flexible, and rapid alternative to conventional recombinant expression systems, capable of producing complex biologics such as enzymes, vaccines, and antibodies. Historically, the recombinant expression of therapeutic peptides in plants has proven difficult, largely due to their small size and instability. However, some plant species harbour the capacity for peptide backbone cyclization, a feature inherent in stable therapeutic peptides. One obstacle to realizing the potential of plant-based therapeutic peptide production is the proteolysis of the precursor before it is matured into its final stabilized form. Here we demonstrate the rational domestication of *Nicotiana benthamiana* within two generations to endow this plant molecular farming host with an expanded repertoire of peptide sequence space. The *in planta* production of molecules including an insecticidal peptide, a prostate cancer therapeutic lead and an orally active analgesic are demonstrated.

## Introduction

Plant-based production of therapeutics offers opportunities for production at scale, with a reduced environmental footprint (Fischer and Buyel, 2020). Although plant-based expression systems are relatively underexplored compared to bacterial or mammalian cell recombinant technologies, they offer capacity for post-translational modification that could enhance or expand the utility of recombinant products. Their green advantage arises from their serum-free production and culture at scale, with only inexpensive inputs required such as fertiliser, water, and light (Nandi et al., 2016; Walwyn et al., 2015). Furthermore, animal cell-free systems negate the threat of zoonotic contaminants, which have derailed product rollouts in the past (Barone et al., 2020). Unlike mammalian or bacterial cell production lines which have the benefit of 50+ years of strain selection (Wuest et al., 2012), plant molecular farming (PMF) is a frontier technology in pharmaceutical production, ripe for genetic and process improvements.

PMF typically employs transient gene expression, which is both rapid and flexible to produce therapeutic antibodies, enzymes, and vaccines. Indeed, transient expression in *Nicotiana* enabled the rapid scale up and deployment of the first antibody therapy against Ebola (Gomez et al., 2021; Olinger et al., 2012). Plant-based production of a seasonal adjustable quadrivalent influenza vaccine is now in phase 3 clinical trials (Ward et al., 2021b), and the development of SARS-CoV-2 vaccines are underway (Ward et al., 2021a). Although some vaccines, enzymes, and antibodies produced in plant-based systems are on the market or in late-stage trials, there remains a gap in terms of peptide production. There is an opportunity to develop plant systems for producing stabilised therapeutic peptides, by capitalising on recent advances in the application of plant peptide ligation machinery (Jackson et al., 2020; Rehm et al., 2021).

Some plants natively produce stable head-to-tail cyclic disulfide-rich peptides (cycDRPs), with the best characterised being the three-disulfide containing cyclotide peptide family (Craik et al., 1999) and the single-disulfide containing sunflower trypsin inhibitor (SFTI) peptide (Luckett et al., 1999). These peptides are gene encoded and processed from larger precursor proteins as they transit through the plant endomembrane system. The final maturation step is predicted to occur in the vacuole where backbone cyclisation is performed by a class of ligase-competent asparaginyl endopeptidases (AEPs), found only in five angiosperm families (Jackson et al., 2018; Jackson et al., 2020). Once extracted, cyclotides are highly stable, and tolerant of a range of thermal, proteolytic or chemical insults (Colgrave and Craik, 2004), making them valuable scaffolds for peptide engineering applications (Wang and Craik, 2018). One proven application is for bioactive epitope grafting, which helps stabilise and configure an epitope for improved efficacy and therapeutic half-life (Chan et al., 2016; Chan et al., 2011). Thus, cycDRPs are envisioned as customisable vehicles to carry therapeutic sequences.

Although grafted cycDRPs can be produced synthetically at laboratory scale, their production at commercial yields is highly suited to a plant-based system, where both precursor and ligase capable AEP can be co-operatively stacked. However, in plants generally most suited as biofactory hosts, such as *Nicotiana benthamiana*, the endogenous AEPs have not evolved for peptide ligation, but rather retain their ancestral function as hydrolases. Thus, these endogenous AEPs may have a negative influence on the resulting cyclic peptide yield, from either out competing and hydrolysing the precursor, or by linearising any cyclic peptide product formed. Once mis-processed, peptides lack the bond energy required for transpeptidation, thus endogenous AEP activity can have a significant negative impact on the sequence space available for PMF of cyclic peptides.

The development of gene editing tools like CRISPR has enabled targeted genetic improvements of crop species, previously thought impossible (Doudna and Charpentier, 2014). Here we demonstrate the genetic customisation of the industrialized PMF crop *Nicotiana benthamiana* through rapid and rational ‘domestication’ enabling the accumulation of heretofore unattainable peptides with therapeutic potential in a plant-based system. We describe our simple and effective genomic edits that enable the production of therapeutics to treat prostate cancer, Netherton syndrome, and neuropathic pain. We further show the production of a potent insecticidal peptide naturally produced in garden pea.

## Results

### Cyclic SFTI-1 therapeutic peptide candidates harbouring Asn residues are inefficiently produced in *N. benthamiana*

SFTI-1 is a 14aa cyclic peptide naturally produced in sunflower seed, and is a favoured scaffold for peptide engineering applications (de Veer et al., 2021). Previously, using *N. benthamiana* as a host for transient gene expression, we demonstrated the successful *in planta* production of SFTI-1 as well as a variant displaying low picomolar inhibition of the human serine protease plasmin (Jackson et al., 2019; Swedberg et al., 2019). In the same study we attempted to improve the cyclisation yield by substituting the Asp residue of SFTI-1 with an Asn, which we predicted, for some AEPs, could improve the cyclisation efficiency. However, unlike the native SFTI-1, SFTI-1_N could not be produced *in planta*, which led to the hypothesis that the Asp-Asn residue exchange is detrimental to the stability of the peptide *in planta*.

We first aimed to determine if this problem of instability is more broadly applicable to other SFTI-1 therapeutic candidates, including examples proposed as potential leads for the treatment of prostate cancer. To approach this question, we assembled three additional expression constructs, two encoding for SFTI-1 variants designed to inhibit human kallikrein 4 (SFTI-1_KLK4_D and SFTI-1_KLK4_N) (Riley et al., 2019; Swedberg et al., 2011) and one encoding for a kallikrein 5 inhibitor (SFTI-1_KLK5_N) (de Veer et al., 2016). The two kallikrein 4 inhibitors were identical, apart from an Asp – Asn exchange at the cyclisation residue. This single residue change (Asp-Asn) was previously demonstrated to confer a 125-fold increase in potency (K_i_ = 0.04 nM) and enhanced selectivity over off-target serine proteases (Swedberg et al., 2011). Thus, of the two peptides, SFTI-1_KLK4_N represents the preferred candidate for plant-based production. The kallikrein 5 inhibitor was likewise chosen as a good test peptide, where the best performing peptide harboured an Asn residue at the AEP processing site and was most potent (K_i_ – 4.2 nM) (de Veer et al., 2016).

For expression *in planta*, we assembled SFTI-1 peptide expression constructs by incorporated back translated SFTI-therapeutic candidates into the Oak1 precursor framework, replacing the sequence encoded the cyclotide kb1 (**Figure 1a**). To ensure high-level transient expression in *N. benthamiana* we assembled each designer precursor peptide gene and AEP ligase gene into the pEAQ vector system, which allowed for co-infiltration (Sainsbury et al., 2009). As previously demonstrated for SFTI-1 (Jackson et al., 2019), infiltration of *N. benthamiana* leaf with Agrobacterium harbouring precursor gene constructs alone produced no detectable cyclic peptide masses as assessed using MALDI-TOF-MS (data not shown). However, upon co-infiltration with Agrobacterium harbouring the expression vector for ligase competent AEP (OaAEP1b), backbone cyclic masses were observed for native SFTI-1 and SFTI-1_KLK4_D (**Figure 1b**). For the remaining three peptides, each containing an Asn at the cyclisation residue, no cyclic masses were evident suggesting that these peptides are highly unstable in *N. benthamiana* leaf cells.

**Fig 1.**
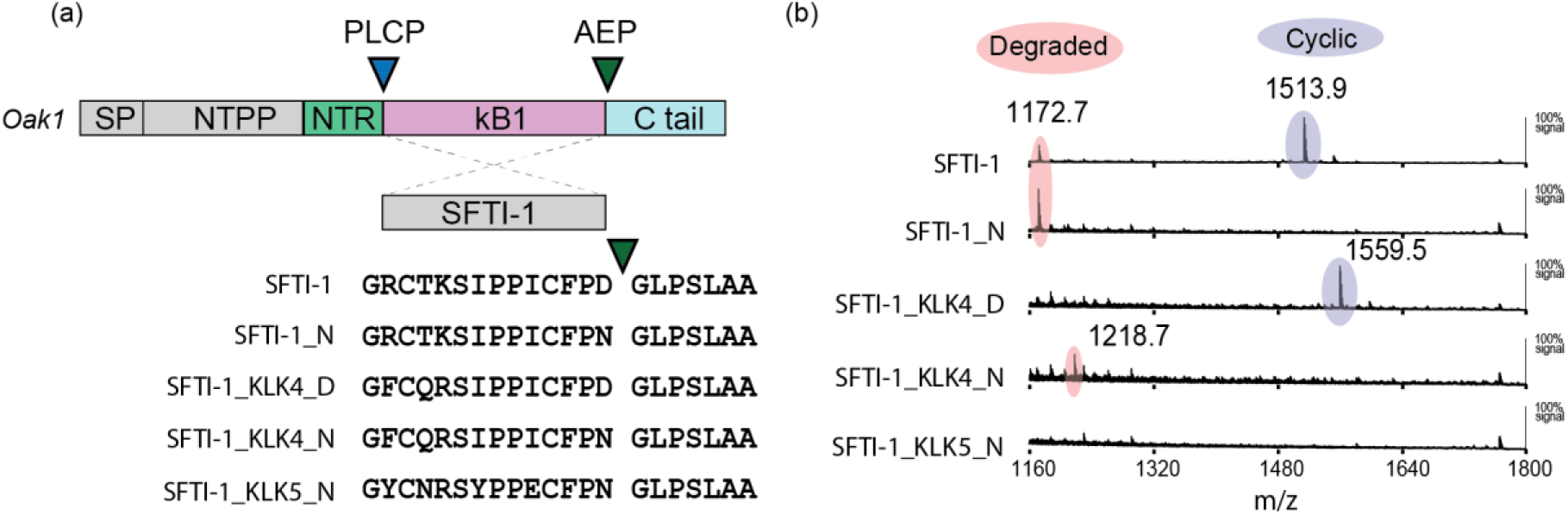
**(a)** SFTI-1, SFTI-1_N and therapeutic peptide candidates were prepared for plant-based expression by insertion of peptide coding sequence into the *Oak1* gene, replacing the sequence domain for the cyclotide kB1. The processing of the engineered SFTI-1 precursors is predicted to be controlled by a papain like cysteine protease (PLCP) and an asparaginyl endopeptidase (AEP) at the N and C-terminus respectively. The Oak1 signal peptide (SP), amino terminal propeptide (NTPP) and amino terminal repeat (NTR) sequence ensures that the precursor enters the endomembrane system with targeting towards the vacuole. **(b)** Co-expression of engineered SFTI-1 precursors with the AEP ligase OaAEP1b in *N. benthamiana* resulted in MALDI-MS detection of either degraded peptides (shaded in peach) or cyclic peptides (shaded in purple).

### A quadruple AEP knockout depletes competing AEP activity with minimal deleterious phenotypic effects

As the minimal change required to induce *in planta* instability of SFTI-1 and the tested variants was a single Asp-Asn substitution, we hypothesised that endogenous AEP activity at Asn residues is problematic. To test this hypothesis, we simultaneously produced lesions in four *N. benthamiana* AEP sequences using CRISPR-Cas9. The chosen loci Niben101Scf04675g08014.1, Niben101Scf04539g04014.1, Niben101Scf18356g00003.1, and Niben101Scf04779g01004.1 were given the gene symbols NbAEP1, NbAEP2, NbAEP3, and NbAEP4 respectively.

AEP knockout lines were produced by introducing an array of four gRNA-tRNA repeats, each targeting one of the selected AEPs, into pKIR1.1, which is a CRISPR/Cas9 expression vector carrying the pFAST seed selection system conferring expression of mRFP in seeds (Tsutsui and Higashiyama, 2016). A hemizygous primary transformant was allowed to self with the resulting seed negatively selected for mRFP expression, thus giving rise to a series (n=7) of genome-edited genotypes in the T_1_ generation that were no longer carrying the pKIR1.1 CRISPR/Cas9 cassette. Sites targeted by the crRNA array have restriction enzyme sites present near the protospacer adjacent motif (PAM) end of the crRNA to enable the use of the site as a cleaved amplified polymorphic sequence (CAPS) marker (**Figure S1**). In this way transgene-free T_2_ genotypes carrying some form of mutation at the target loci were identified.

Seven plant lines were selected from the T_2_ population for further analysis. Amplicons from gDNA containing target sites were examined by Sanger sequencing to categorize mutations present. The lesions identified in the T_2_ led to non-silent point mutations, frameshifts and premature stop codons at the loci targeted. However, some combination of bi-allelic states were observed for all NbAEPs (data not shown) in the population of selected plants, highlighting the need to select for a quadruple homozygous individual. Plants were allowed to self and quadruple mutants homozygous at the chosen loci were identified in the T_4_ generation by CAPS screening and again by validating allelic states with Sanger sequencing of amplicons (**Figure S1**). In this way a single line, harbouring mutations in all four targeted AEPs was selected and named ΔAEP. The ΔAEP genotype is homozygous for an 18 base pair deletion encompassing the splice acceptor site of exon 4 in NbAEP1, insertion of a single nucleotide at position 388 of the NbAEP2 coding sequence, a single nucleotide insertion at position 544 of the NbAEP3 coding sequence, and a missense mutation at position 538 of the NbAEP4 coding sequence.

RNA-seq analysis was performed on T_5_ ΔAEP and wild-type plants to further validate the structure of the resulting NbAEP transcripts and observe global gene expression changes between genotypes in a transient expression experiment. Both ΔAEP and wild-type plants were infiltrated with pEAQ-eGFP and samples of infiltrated tissue taken for RNA-seq (n=2 per genotype). Reads of ΔAEP were mapped against the Sol Genomics *N. benthamiana* Niben101 transcript set and visualized to validate changes in NbAEP mRNA, particularly at NbAEP1. The ΔAEP NbAEP1 allele produced transcripts with disrupted exon splicing at exon 4 as seen by a lack of read coverage at the crRNA target site (**Figure S1**), thus confirming the NbAEP1 allele present in ΔAEP as null. Expression levels were estimated and revealed that NbAEP1, NbAEP2, and NbAEP3 were significantly down-regulated in ΔAEP (>2.9-fold down-regulated, p-values <0.007) (**Table S1**). NbAEP4 exhibited very low expression in both genotypes such that it did not result in mapped reads above the threshold for abundance estimation. In addition to the downregulated AEP transcripts in ΔAEP, we observed one other transcript that was significantly downregulated less than 2-fold, and 13 transcripts significantly upregulated more than 2-fold (**Table S1**). In summary, our ΔAEP genotype resulted in minimally significant perturbations to the Agrobacterium-infiltrated wild type *N. benthamiana* transcriptome. Furthermore, ΔAEP plants were morphologically indistinguishable from wild type. A simple biomass experiment using a randomized plot design under artificial light and growth conditions failed to reveal significant differences in biomass between genotypes (mean DW / 4-week-old plant (n=10): wild-type 4.256g ±0.371g, ΔAEP 4.281g, ±0.665g).

To determine the effectiveness of our AEP knockout strategy in transient expression of peptides, we first assessed and compared processing of a modified cyclotide precursor gene *Oak1_HIIAA* in wild type and ΔAEP plants (**Figure 2**). It was demonstrated previously that *Oak1* expression in *N. benthamiana* without co-expression of a helper AEP ligase results in accumulation of linear, linear extended, linear truncated, and a small MS signal representing cyclic kB1 (Gillon et al., 2008; Saska et al., 2007). Of these, only the cyclic, full length linear and linear minus a Gly at the N-terminus can be attributed to AEP processing, with the remainder representing processing by carboxypeptidases that compete for the substrate (**Figure 2A**). For our analysis, to ensure that MALDI-TOF-MS could be used to differentiate all predicted processed products we modified the CTPP of Oak1 to HIIAA from the natural GLPSLAA, as with the later construct it is impossible to differentiate the linear kB1 mass from a linear form harbouring a truncated N-terminal Gly and C terminal extended Gly. Expression of *Oak1_HIIAA* in wild type *N. benthamiana* resulted in MS signals (as a percentage of total kB1 related signals) of 7.6 +-2.7% for cyclic, 20.1 +-1.8% for linear and 16.4 +-6.4% for linear – Gly, with the remaining ∼58% of signal being for peptides carrying C terminal extensions with or without the N-terminal truncation event. In contrast, expression within the ΔAEP line produced almost no detectable signal for cyclic, linear, or linear – Gly peptide, with ∼100% of the signal representing non-AEP processed forms (**Figure 2B, C**). These results clearly demonstrate the effectiveness of our AEP knockout strategy to remove interfering endogenous AEP activity.

**Fig 2.**
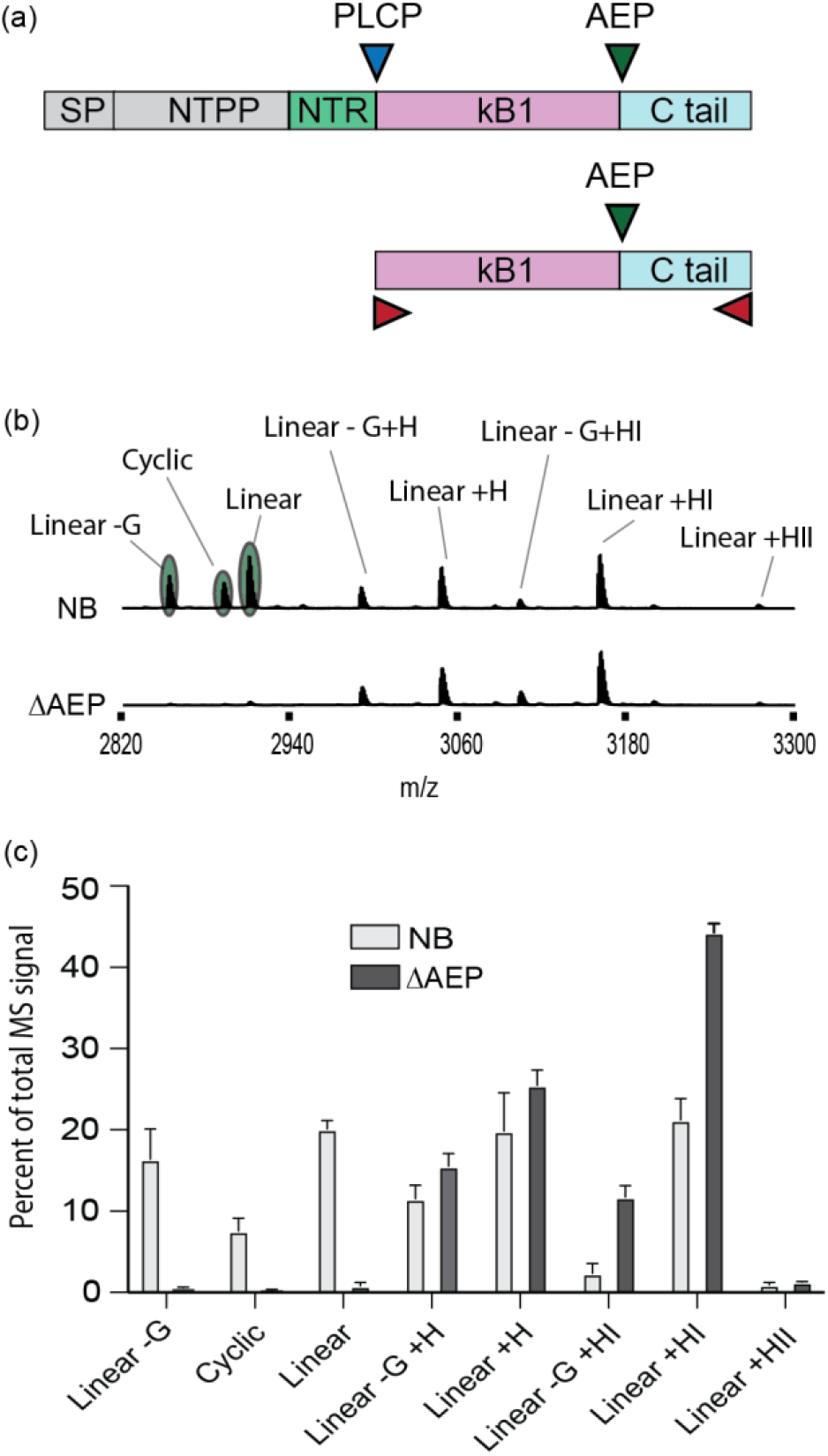
**(a)** The Oak1 precursor is predicted to be processed initially at the N-terminus by a papain like cysteine protease (PLCP) (blue triangle), followed by asparaginyl endopeptidase (AEP) processing (green triangle) at the C-terminus. Without the co-expression of an AEP ligase, endogenous AEPs that prefer hydrolysis over ligation will compete for the expressed substrate with amino and carboxypeptidases (red triangles). **(b)** MALDI-TOF-MS analysis (representative) of *Oak1_HIIAA* expression in wild type *N. benthamiana* (NB) alongside the ΔAEP gene edited accession. Only the signals highlighted in green can be attributed to endogenous AEP processing with the remainder representing C-terminal processing of the precursors C-terminus. **(c)** Mean and standard deviation (n=3) of the individual kB1 identified masses as a percent of the total kB1 peptide MS signal detected by MALDI_TOF-MS.

### AEP depletion expands accessible peptide sequence space enabling the expression of cyclic therapeutic peptide leads

Having established a *N. benthamiana* plant line with reduced endogenous AEP activity, we next tested our hypothesis that reduced endogenous AEP activity would have a positive influence on the yield of Asn-containing SFTI-1_KLK4_N and SFTI-1_KLK5_N. We repeated infiltration experiments and compared relative yields obtained from wild type and ΔAEP plants. Like previously demonstrated, close to no MS signal for cyclic SFTI-1_KLK4_N and SFTI-1_KLK5_N could be detected in infiltrated wild type plants. In contrast, MS signals for cyclic SFTI-1_KLK4_N and SFTI-1_KLK5_N were readily detectable when constructs were infiltrated into the ΔAEP genotype (**Figure 3A, B**), representing a considerable breakthrough. For the expression of cyclotides kB1 and kB2, relative yields were slightly reduced in ΔAEP plants. A subsequent test of another AEP ligase CtAEP1 in ΔAEP plants gave a similar result (**Figure S2**). Further clarification of the negative effect in the ΔAEP genotype for kB1 and kB2 substrates is planned with a reduced transactivation of AEP ligase likely playing a role.

**Fig 3.**
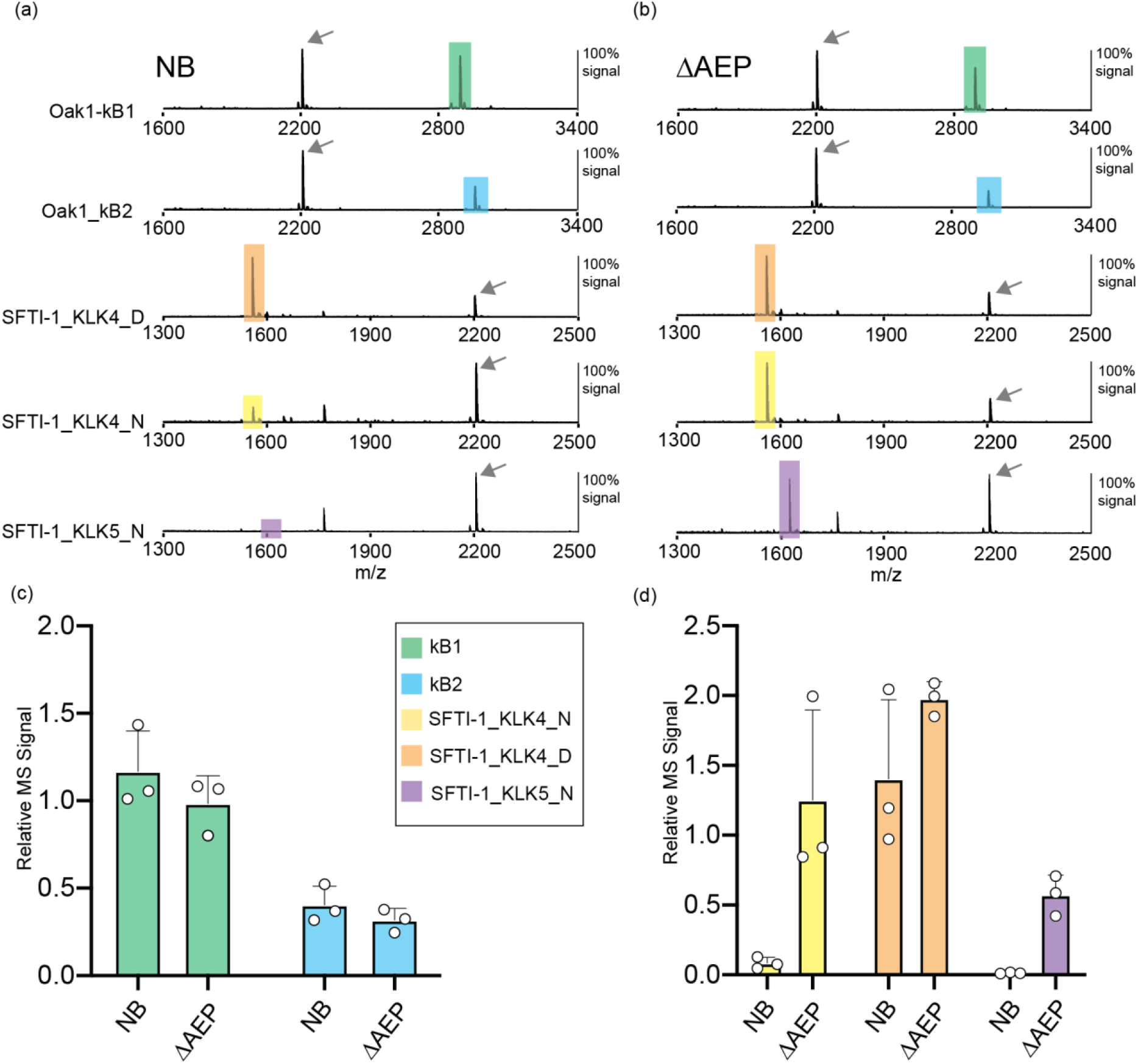
Representative MALDI-TOF-MS of cyclic peptides accumulated in **(a)** wild type *N. benthamiana* (NB) and **(b)** the ΔAEP accession. MS signals for cyclic peptides are highlighted to match the colours in panels c and d. An arrow indicates the MS signal for the internally spiked peptide control that served to normalise MS signals for relative quantification. **(c)** Mean and SD (n=3) of relative kB1 and kB2 MS signals detected in crude peptide extracts of infiltrated *N. benthamiana* (NB) and the ΔAEP accession. **(d)** Mean and SD (n=3) of relative SFTI-1_KLK4_N, SFTI-1_KLK4_D and SFTI-1_KLK5_N MS signals detected in crude peptide extracts of infiltrated *N. benthamiana* (NB) and the ΔAEP accession.

### Cyclotides are more resistant than SFTI-1 molecules to endogenous AEP activity

In common with all plant species, cyclotide producers encode AEPs as a multi-gene family with isoforms differing in their propensity for peptide ligation (Harris et al., 2019; Serra et al., 2016). Thus cyclotides, which naturally co-locate with AEPs in vegetative cell vacuoles (Conlan et al., 2011; Slazak et al., 2016), may have evolved structures that are more resistant to hydrolytic AEPs than for the Asn containing SFTI-1 peptides tested in this study. To gain further insight into this we set up an infiltration experiment in our ΔAEP plant where we co-expressed AEP ligase (OaAEP1b), individual NbAEPs and the precursor genes of Oak1 or SFTI-1_KLK4_N (**Figure 4**). For SFTI-1_KLK4_N, co-expression of any of the four tested *N. benthamiana* AEPs resulted in a complete elimination of cyclic product formation, as is observed in wild type plants (**Figure 4A**). In contrast, Oak1 processing appears to be only moderately affected with cyclic kB1 predominating the MS profile, irrespective of NbAEP overexpression (**Figure 4B, C**).

**Fig 4.**
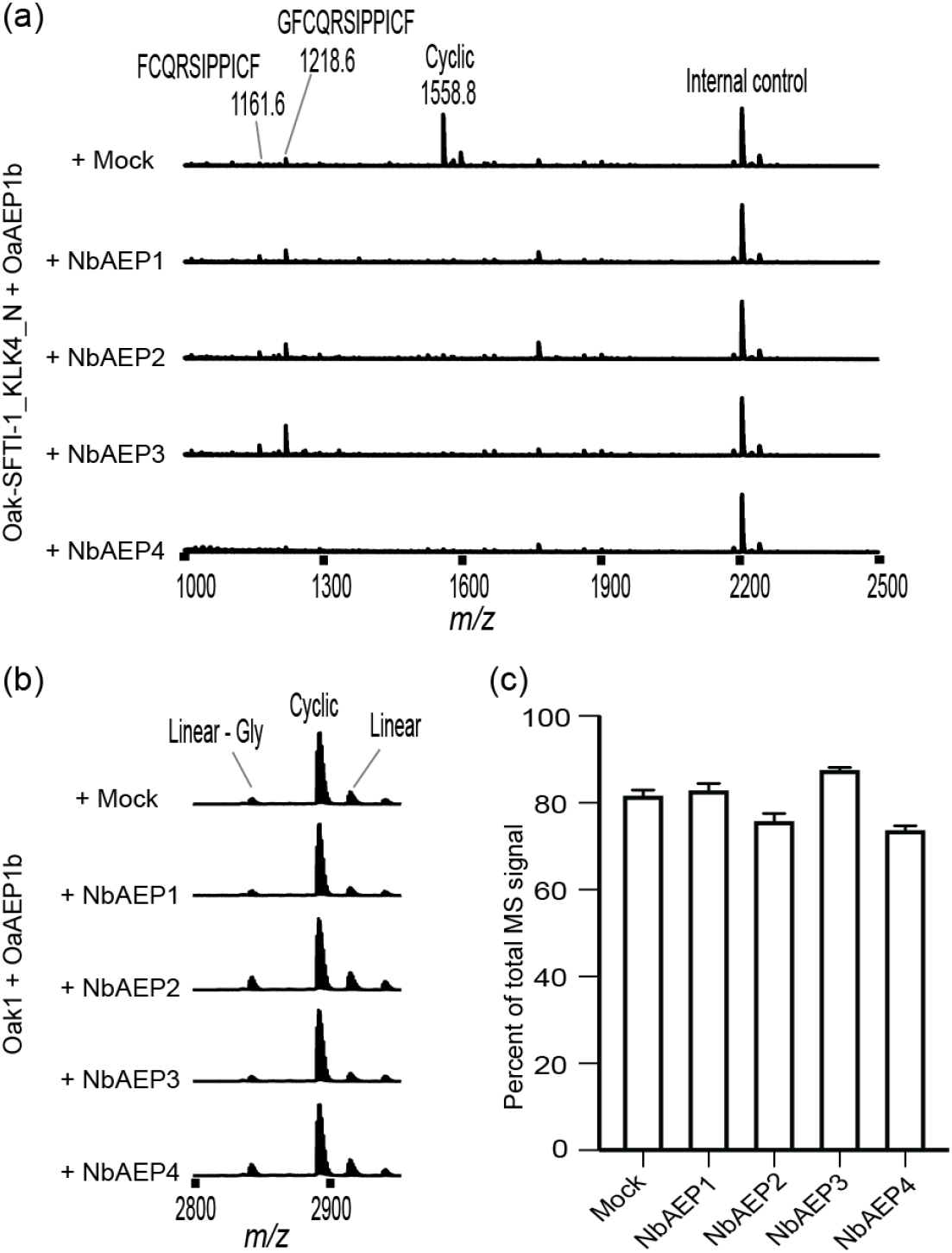
Representative MALDI-TOF-MS of **(a)** cyclic SFTI-1_KLK4_N and **(b)** cyclic kB1 accumulation in the ΔAEP accession upon co-expression of peptide precursor, AEP ligase and NbAEP genes. **(c)** Mean and SD (n=3) of the percent of MS signal representing cyclic kB1 upon co-expression.

### Bioactive linear peptide accumulation

Having shown that we could improve the *in planta* yield of cyclic SFTI-1 therapeutic leads, we next aimed to determine if the same could be true for bioactive peptides harbouring internal Asn sites. We first chose to test expression of the pea albumin-1 gene (*PA1*) that encodes for the 37aa disulfide rich insecticidal peptide Pa1b (Higgins et al., 1986). This peptide is of high interest for development as it represents the first ever peptide that specifically inhibits insect vacuolar proton pumps (Chouabe et al., 2011). Although produced naturally in many legumes, the development of Pa1b as a commercial insecticide would benefit from an expression platform allowing the rapid testing of variants for improved yield and potency, and with the capacity for scaled up production (Eyraud et al., 2013). Pa1b has two internal Asn sites that represent putative processing sites of endogenous AEPs, thus Pa1b represents a good peptide to test for yield improvements in our ΔAEP *N. benthamiana* line devoid of AEP activity (**Figure 5A**). Similar to the *PA1* expression results reported by Eyraud et al. (2013), we found that *PA1* expression in *N. benthamiana* results in a number of processed forms, with the predominant signals representing Pa1b with the C-terminal Gly37 removed, with or without oxidation (plus 16 daltons) of the methionine residue. We found this similar pattern of processing in both wild type and ΔAEP plants; however relative yields were calculated to be ∼3.7-fold and ∼1.9-fold higher in ΔAEP plants for Pa1b-Gly and Pa1b-Gly+Met^ox^ respectively (**Figure 5B and C**). By scaling up plant infiltration, Pa1b yield was calculated at ∼0.2 mg/ g DW tissue at 95% purity, and the plant derived peptide was shown to be cytotoxic to Sf9 cells with a CC50 of 13.58nM (**Figure S3**).

**Fig 5.**
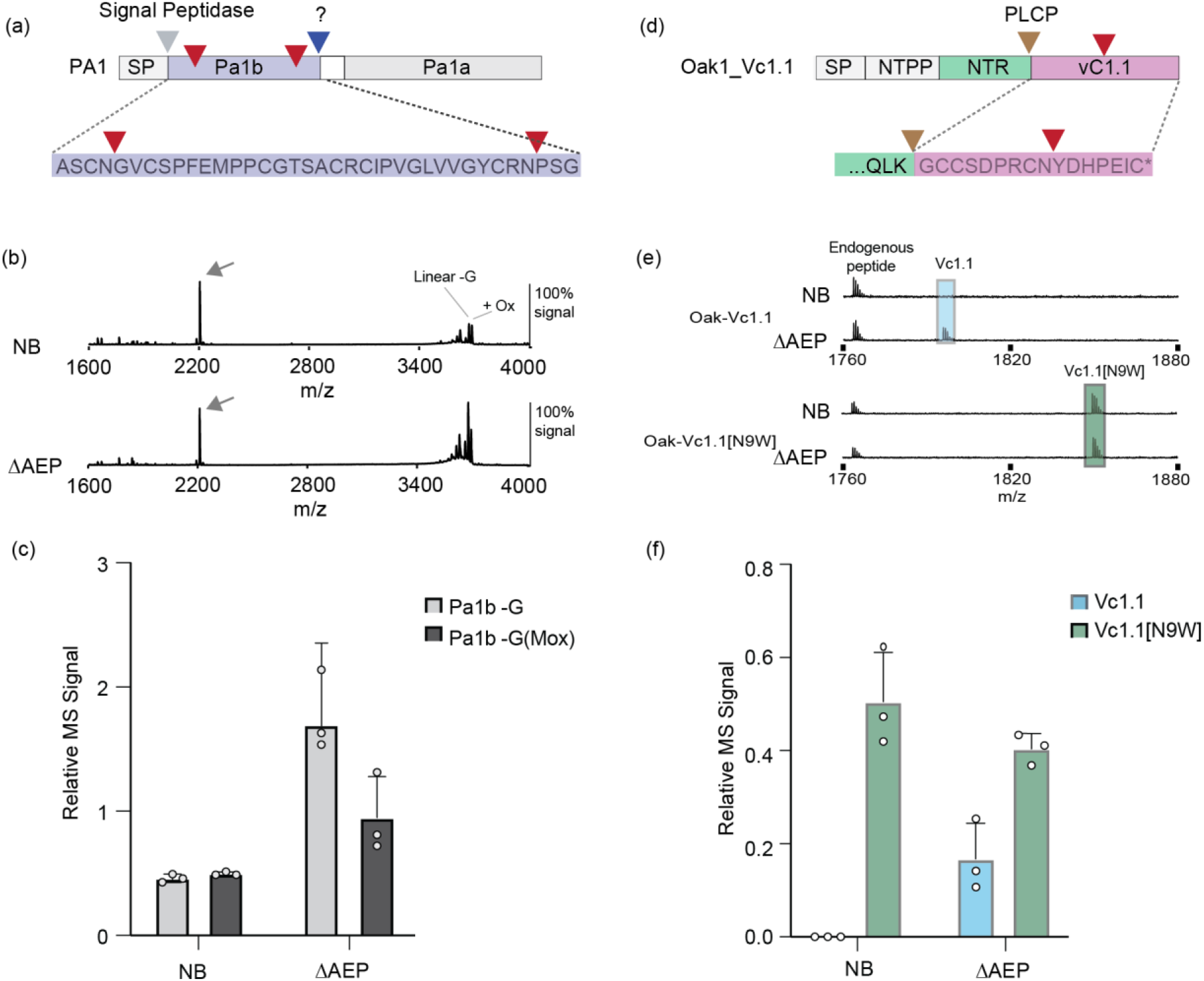
**(a)** The insecticidal peptide Pa1b is encoded by the pea albumin 1 gene (*PA1* gene) that additionally encodes for the larger PA1a domain of unknown function. The Pa1b N-terminus is predicted to be released via a signal peptide (SP) cleavage event during co-translation into the endoplasmic reticulum. The protease responsible for the release of the Pa1b C-terminus is unknown. Two internal Asn sites are present (red triangles) that may represent putative AEP processing sites. **(b)** Representative MALDI-TOF-MS of crude peptide extracts prepared from infiltrated leaves of *N. benthamiana* (NB) and ΔAEP. Indicated by an arrow is the MS signal for the internally spiked peptide control that served to normalise MS signals for relative quantification. **(c)** Mean and SD (n=3) of the relative MS signal representing Pa1b-G and Pa1b-G(Mox) peptides. **(d)** The conotoxin Vc1.1 and its analogue Vc1.1[N9W] were prepared for plant-based expression by insertion of peptide coding sequence into the *Oak1* gene, replacing both the cyclotide kB1 domain and C-terminal tail. Processing was predicted to be controlled by a papain like cysteine protease (PLCP) (brown arrow). One internal Asn site (indicated by a red arrow) is present in Vc1.1, substituted for a Trp in Vc1.1[N9W]. **(e)** Representative MALDI-TOF-MS of crude peptide extracts prepared from infiltrated leaves of *N. benthamiana* (NB) and ΔAEP **(f)** Mean and SD (n=3) of the relative MS signal representing Vc1.1 and Vc1.1[N9W].

As an example of a non-plant peptide with therapeutic potential, we prepared constructs (**Figure 5D**) for the expression of the 16 aa two disulfide containing conotoxin Vc1.1, which is a lead compound for the treatment of neuropathic pain (Clark et al., 2010). Native Vc1.1 harbours one internal Asn site but structure activity studies have identified that this residue can be substituted with a tryptophan with minimal changes to activity (Yu et al., 2013). This allowed us to test and compare the expression of the Asn containing Vc1.1 and Vc1.1[N9W] in wild type plants and in our ΔAEP line. Of the two variants, only Vc1.1[N9W] could be detected in wild type plants, indicating that this Asn processing site is liable for processing (**Figure 5E, F**). In our ΔAEP line, both Vc1.1 and Vc1.1[N9W] could be detected, suggesting that reducing endogenous AEP levels is beneficial for the accumulation of linear Vc1.1, in a similar fashion to Pa1b.

## Discussion

The uptake of *N. benthamiana* for industrial plant molecular farming is gaining momentum, with commercial entities employing *N. benthamiana* as the preferred biofactory host in Africa, North America, and Europe. In particular, the ‘Lab’ strain, derived from a wild *N. benthamiana* accession, which naturally carries a genetic lesion that reduces transgene silencing, has been the dominate accession for transient-expression based production for PMF (Bally et al., 2018). Genetic customization reported to date of the ‘Lab’ accession has been limited to the alteration of the glycosylation pathway, to humanise recombinant glycoproteins (Jansing et al., 2019). Altering *N. benthamiana* to create new genotypes exhibiting expanded expression capability represents great untapped potential. The work presented here demonstrates that potential, made possibly by multiplex CRISPR gene editing, whereby knocking out four endogenous AEPs enabled the production of previously non-producible peptide products.

The *N. benthamiana* ΔAEP genotype is similar to the quadruple AEP mutant described from Arabidopsis in that it does not exhibit a deleterious phenotype in controlled growth environment settings (Gruis et al., 2004). In Arabidopsis, the quadruple AEP knockout genotype is however impaired in programmed cell death and in the hypersensitive response when challenged with *Pseudomonas syringae*, indicating a role of AEPs in pathogen response (Rojo et al., 2004). Further roles for AEPs include storage protein processing (Shimada et al., 2003), and separation of the endosperm from the testa during seed development, demonstrated in Arabidopsis (Nakaune et al., 2005). In our *N. benthamiana* ΔAEP genotype no deleterious phenotypes were observed in controlled growth conditions and biomass was unaffected. In field conditions or in controlled environments where pathogen infection may occur the ΔAEP genotype might however exhibit greater susceptibility and further exploration of pathogen susceptibility due to reduced AEP function is warranted.

The proteolytic stability of recombinant proteins produced in plants remains a significant challenge. It is clear is that there is not a one size fits all approach to improving stability, with larger recombinant proteins possibly degraded by numerous proteases *in planta* or during the extraction phase. Strategies developed to counter proteolytic interference have included the co-expression of protease inhibitors (Grosse-Holz et al., 2018; Ma et al., 2021), the re-engineering of protease susceptible residues and in directing subcellular compartmentation to protein friendly organelles (Streatfield, 2007). In the case of therapeutic peptide production in plants, our approach for enhancing recombinant peptide stability involves co-expression of a ligase capable AEP with a peptide substrate amenable to AEP mediated backbone cyclisation (Poon et al., 2018). With their lack of termini, backbone cyclic peptides have been demonstrated to have improved stability, both in human serum stability assays (Ganesan et al., 2021; Muratspahić et al., 2021; Wong et al., 2012) and during accumulation in plant cells (Jackson et al., 2019). Ligase type AEPs, essential for backbone cyclisation, however, have only been identified from cyclotide producing plant species (Jackson et al., 2020), thus they must be exogenously co-expressed in biofactory hosts such as *N. benthamiana*.

Through CRISPR-Cas9 gene editing, we have shown here that the reduction of endogenous AEP activity in *N. benthamiana* positively influences the accumulation of the Asn containing cyclic SFTI-1 therapeutic peptide candidates (SFTI-1_KLK4_N (Riley et al., 2019; Swedberg et al., 2011) and SFTI-1_KLK5_N (de Veer et al., 2016)). This interference by endogenous AEPs appears particularly problematic for Asn containing SFTI-1 molecules, and not for cyclotides, which have likely evolved to remain resistant to AEP-mediated hydrolysis. Mechanistically, Asn-containing SFTI-1 molecule breakdown may occur through either endogenous AEPs outcompeting the transgene derived AEP ligase, or by endogenous AEP activity re-cleaving any cyclic peptide generated. We further demonstrate improved production levels upon transient expression in the ΔAEP genotype of the linear peptides vc1.1 and Pa1b, that each carry internal Asn residues. Of particular interest was the 1.9-to-3.7-fold increase in the relative yield of Pa1b-Gly+Met^ox^ and Pa1b-Gly respectively when compared to expression in wild type *N. benthamiana*. Interestingly using MALDI-TOF-MS, no intermediate Asn specific cleavage products were detectable, suggesting rapid hydrolysis of peptide fragments after the primary AEP cleavage event. Thus, this inefficiency in Pa1b production in wild type *N. benthamiana* could not be forecast, suggesting that many other small Asn containing peptides could unknowingly benefit from production in the ΔAEP genotype. Similarly, when the vc1.1 peptide was expressed in wild type *N. benthamiana*, no peptide at all could be detected, which could be attributed to either protease-mediated instability or inefficient peptide folding. Only by expressing in the ΔAEP genotype could we identify that AEP hydrolysis was limiting the accumulation of the vc1.1 peptide. Likewise, other non-accumulating PMF products containing internal protease sites may benefit from select knockout of offending proteases.

## Conclusion

Here we have demonstrated genome editing as a valuable strategy to enable the accumulation of cyclic bioactive peptides in the framework of PMF. In reducing endogenous AEP activity of *N. benthamiana* we have demonstrated an expansion of the repertoire of peptides that this favoured biofactory plant can achieve. The targeting of interfering proteases is an approach that may be applied to other recombinant products that may have failed previously. The work thus potentially has broad implications for the efficient and economical recombinant production of therapeutic and insecticidal peptides.

## Experimental procedures

### Generation of ΔAEP *N. benthamiana* genotype

An AEP knockout line was produced by introducing an array of four gRNA-tRNA repeats, each targeting one of the selected AEPs, into pKIR1.1 which is a CRISPR/Cas9 expression vector carrying the pFAST seed selection system conferring expression of mRFP in seeds (Tsutsui and Higashiyama, 2016). In pKIR1.1, the crRNA is cloned into the relevant site using AarI, a type IIS restriction enzyme that liberates four bases of unique sequence on both sides of the plasmid backbone. Arrays of tandem gRNA-tRNA pairs, each targeting a single NbAEP gene, were constructed by assembly of tiled PCR products (**Supplementary Methods**). Tiles were assembled into a modified pGEM-T plasmid backbone using the NEBuilder HiFi kit. The completed gRNA-tRNA array was digested using AarI and ligated into AarI-digested pKIR1.1 using NEB T4 ligase as per manufacturer’s protocol.

To enable transformation of *N. benthamiana* the pKIR1.1_ ΔAEP construct was transferred into *Agrobacterium tumefaciens* (strain LBA4404) by electroporation. Transformation of *N. benthamiana* leaf discs and regeneration of planets was performed as described by (Clemente, 2006) with 30mg/L hygromycin B used for selection (Pavli et al., 2011). Shoots were rooted in rooting medium as described by Clemente (2006) supplemented with 10mg/L hygromycin B. Primary transgenics were acclimatised in a controlled environment room and grown to maturity where progeny seed was scored for segregation of red fluorescence using a Nikon SMZ18 stereo microscope equipped with a mercury lamp and mRFP filter set. Non-fluorescent seed, devoid of Cas9 expression, were picked up with a moistened 27-gauge needle, stored in a 1.5mL tube, and when ready for planting, grown to maturity. Progeny were screened for lesions at AEP loci using CAPS markers and Sanger sequencing (**Supplementary Methods**). Amplicons for gDNA fragments surrounding gRNA target sites were digested, with those amplicons hosting polymorphisms remaining undigested and those matching wild-type alleles visualized as digested on agarose gels. CAPS positive plants were then selected for Sanger sequencing. A single line (termed ΔAEP) was selected for use in all infiltration experiments.

### Plant cultivation and material

*N. benthamiana* and ΔAEP plants were cultivated using a hydroponic nutrient system in a controlled plant growth facility as part of the Clive and Vera Ramaciotti Facility for Producing Pharmaceuticals in Plants. Temperature in the growth room was set at 28°C and plants were grown under 170µmol m-2 s-1 of LED illumination in 16 hr daylight. (AP67 LED spectra, Valoya Oy, Finland).

### RNA-seq analysis

To determine the global gene expression changes between ΔAEP and wild-type plants, two replicates of each plant were vacuum infiltrated with *Agrobacterium tumefaciens* carrying pEAQ-eGFP. After 4 days, the plant tissue was sampled, ground to powder in liquid nitrogen and RNA isolated using TRIzol reagent (ThermoFisher Scientific) using the standard manufacturer’s protocol. Samples were treated with RNAse-free DNAse (Ambion), quantified by spectrophotometry, visualized on an agarose gel to check integrity, and submitted for Illumina NextSeq 500 RNA-seq for mRNA input (Australian Genome Research Facility, AGRF). Data are available from the National Center for Biotechnological Information Sequence Read Archive in BioProject PRJNA784697. Reads were trimmed and filtered using Trimmomatic (Bolger et al., 2014), mapped to the Sol Genomics *N. benthamiana* transcript set using Bowtie2 (Langmead and Salzberg, 2012), quantified with kallisto (Bray et al., 2016), and differential gene analysis performed by EBSeq (Leng et al., 2013). Significantly upregulated or downregulated genes were identified by a cut off p-value of 0.05 and differential genes further refined by a ± 2-fold difference between wild-type and ΔAEP.

### Transient expression in *Nicotiana benthamiana* leaf

Peptide expression constructs were ordered as gene blocks (**Figure S4**) (Integrated DNA Technologies, IDT) with codon usage optimised to *N. benthamiana* according to IDT optimisation software. Gateway cloning enzymes (BP and LR Clonase II, Invitrogen) were used to clone precursor coding sequence initially into the intermediate pDS221 vector (Du et al., 2020), followed by the plant expression vector pEAQ-DEST1 (Sainsbury et al., 2009). For AEP ligase expression, pEAQ-OaAEP1b and pEAQ-CtAEP1 were used and have been previously described (Jackson et al., 2018). For the expression of NbAEPs 1-4 coding sequences were amplified from prepared *N. benthamiana* leaf cDNA and subsequently transferred to pDS221 and to the plant expression vector pEAQ-DEST1 (Sainsbury et al., 2009). All vectors were validated by Sanger sequencing before transfer to *Agrobacterium tumefaciens* strain LBA4404 for leaf infiltration experiments.

To compare relative peptide yields between *N. benthamiana* and ΔAEP, plants of five to six weeks of age were vacuum infiltrated with Agrobacterium solutions containing equal amounts of AEP and precursor peptide as estimated by mixing cultures for a final OD600 of 0.5for each component. Before infiltration cultures were resuspended in infiltration buffer (10mM MES (2-[*N*-morpholino]ethanesulfonic acid) pH 5.6, 10mM MgCl_2_, 100µM acetosyringone) and allowed to rest for up to 2 hours to induce virulence. Plants were harvested 5 days post infiltration and immediately freeze dried for storage and peptide extraction.

### Peptide analysis and relative quantification

Freeze dried tissue was ground to a fine powder using a GenoGrinder (SPEX SamplePrep). Peptides were extracted using a (50% (v/v) acetonitrile, 1% (v/v) formic acid) solution at a ratio of 20 µL per mg of DW tissue. Peptides were extracted overnight with gentle mixing before centrifugation to pellet insoluble material. 10 µL of the peptide containing supernatant was then mixed with an equal volume of an unrelated control peptide (GCCSDPRCNYDHPEICGGAAGN) and a further 80 µL of an (80% (v/v) acetonitrile, 1% (v/v) formic acid) solution. For MALDI-TOF-MS this diluted and spiked peptide mix was mixed 1:1 with a α-cyano-4-hydroxycinnamic acid (5 mg mL^-1^ in 50% acetonitrile/0.1% TFA/5 mM (NH_4_)H_2_PO_4_) solution before being spotted and dried on a MALDI plate. MALDI-TOF-MS spectra data was acquired using a 5800 MALDI-TOF MS (AB SCIEX, Canada) operated in reflector positive ion mode. For relative yield isotope cluster area corresponding to the peptide of interest normalised to that obtained for the internally spiked peptide control.

### Scale up production of Pa1b and purification

To produce and purify *N. benthamiana* derived Pa1b peptide for cytotoxicity assessment, ten ΔAEP plants were infiltrated by submerging whole plants in Agrobacterium solution and applying and releasing vacuum. Five days post infiltration, plant leaves were sampled, freeze dried and ground to fine powder using a GenoGrinder. Peptides were extracted overnight using 50% (v/v) acetonitrile, 1% (v/v) Formic acid at a ratio of 50 µL per mg of DW tissue. After centrifugation to pellet insoluble material, the peptide containing supernatant was again freeze dried and resuspended in a 10% (v/v) acetonitrile, 1% (v/v) Formic acid before Solid-Phase Extraction (SPE) using a Phenomenex Strata C18-E SPE cartridge with 10g resin capacity. The eluted 10-50% (v/v) acetonitrile, 1% (v/v) Formic acid fraction was then pooled, lyophilised, and reconstituted in 10% (v/v) acetonitrile, 0.1% (v/v) trifluoroacetic acid in preparation for HPLC on a semipreparative Phenomenex Jupiter C18 RP-HPLC column (250 mm x 10 mm, 5 µm particle size) connected to a Shimadzu LC-20AT pump system (Shimadzu Prominence). The purity of peptides was checked by MALDI-TOF-MS and analytic HPLC using a C18 column (Phenomenex, Jupiter® 5 µm, 300 Å, 150 × 2.0 mm).

### Cytotoxicity assessment of recombinantly produced Pa1b

*Spodoptera frugiperda* (Sf9) insect cells were cultured in ESF 921 medium at 27°C, 5% CO_2_ in 75 cm^2^ flasks until 80% confluent. Cells were plated into 96-well tissue culture plates (100 mL, 10,000 cells/well) and grown for 24 h prior to treatment. Peptides were added in triplicate with final concentrations ranging from 0.1 mM to 50 mM, while control wells were incubated with 1% Triton X-100 and medium only. The cells were further incubated for 24 h at 27°C, 5% CO_2_. Ten mL of MTT (3-(4,5-dimethylthiazol-2-yl)-2,5-diphenyltetrazolium bromide, 5 mg/mL in PBS) was added to each well to a final concentration of 0.5 mg/mL and were incubated for 3 h at 27°C, 5% CO_2_. Supernatants were removed, and the insoluble formazan crystals were resuspended in 100 mL of DMSO. The plate was shaken at room temperature for 10 min to fully dissolve the formazan crystals, and the absorbance of the solutions was measured at 600 nm on a Tecan Infinite M1000Pro plate reader.

### Statistical analysis

One-way ANOVA was performed using GraphPad Prism version 9.00 for Mac OS (GraphPad Software, La Jolla California USA).

## Accession numbers

PRJNA784697

## Acknowledgements

This work was supported by grants from the Australian Research Council (ARC) Discovery Program (DP150100443 and DP200101299) awarded to DJC, Thomas Durek, and EKG. DJC was an ARC Australian Laureate Fellow (FL150100146) during the period of this work. This work was supported by access to the facilities of the ARC Centre of Excellence for Innovations in Peptide and Protein Science (CE200100012).

## Contributions

MJ designed and assembled plant expression constructs and performed and analysed plant infiltration experiments. EKG prepared the CRISPR-Cas9 AEP knockout construct, transformed, genotyped, and performed global differential gene expression analysis of the ΔAEP line. LYC and MDH purified the Pa1b peptide and conducted purity checks and cytotoxicity assays. DJC, MJ and EKG planned the experiments.

## Short legends for supporting information

**Figure S1**. crRNA target sites for NbAEP1, 2, 3 and 4, CAPS marker sites, and resulting mutations for the ΔAEP genotype.

**Figure S2**. CtAEP1 (butelase-1) activity assessment in the ΔAEP accession.

**Figure S3**. Purity and cytotoxicity assessment of Pa1b.

**Figure S4**. Gene sequences ordered as dsDNA gene blocks used in this study.

**Table S1**. Genes significantly upregulated and downregulated greater than 2-fold.

**Supplementary Method 1**. Assembly of plant CRISPR Cas9 expression cassette.

**Supplementary Method 2**. Genotyping genome-edited lines using CAPS markers and Sanger sequencing.

