## Supplementary Information for "Rational domestication of a plant-based recombinant expression system expands its biosynthetic range"

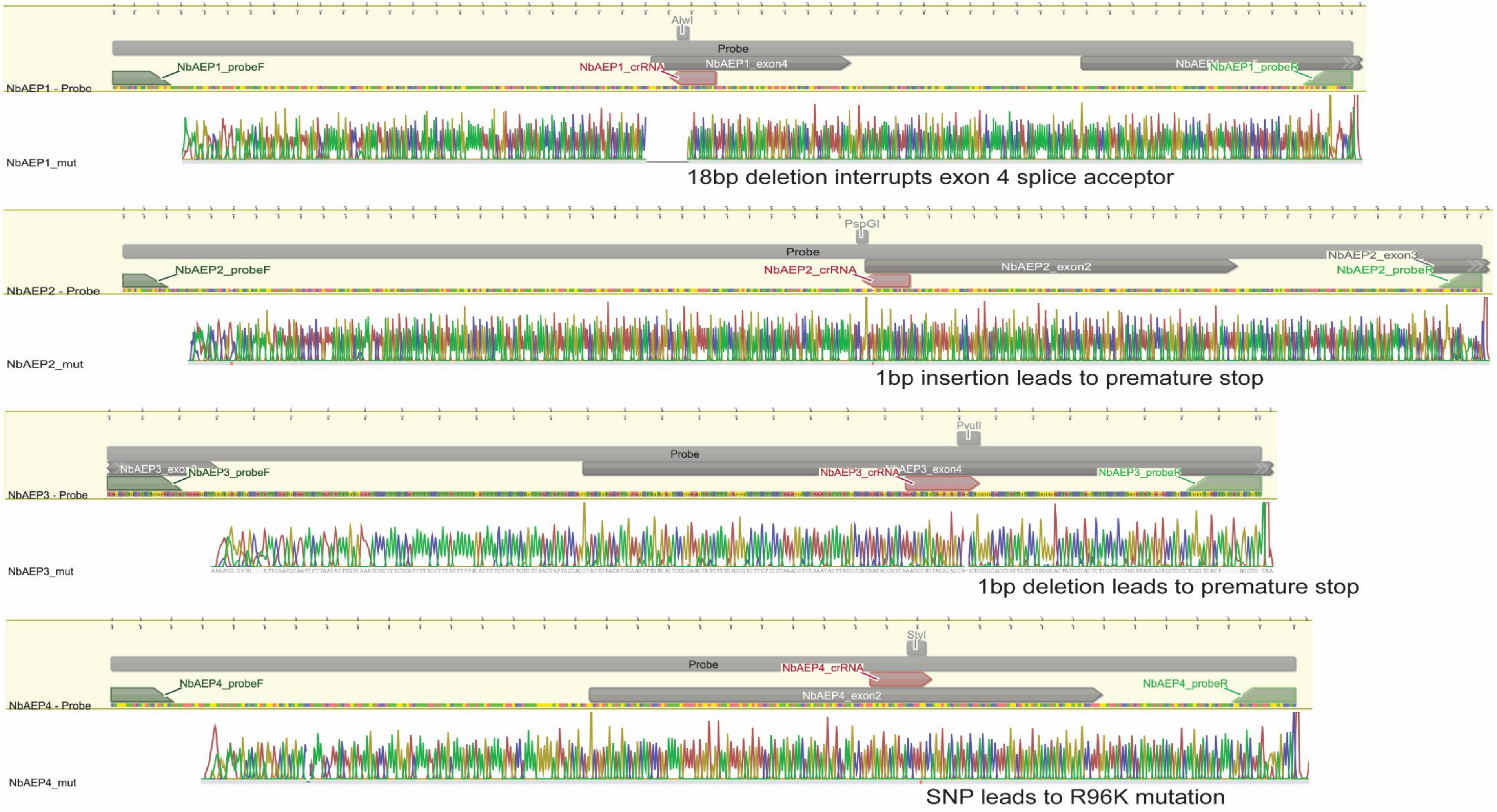


**Figure S1.** crRNA target sites for NbAEP1, 2, 3 and 4, CAPS marker sites, and resulting mutations for the ΔAEP genotype. NbAEP amplicons are given as grey bars labelled as probe, amplified with probeF (forward) and probeR (reverse) primers, and digested using sites annotated by grey rectangles for CAPS marker analysis. Red font and grey and red shapes denote the location of crRNA targets. Notation on the allelic change from wild-type is given under the representative Sanger sequencing trace for each locus.

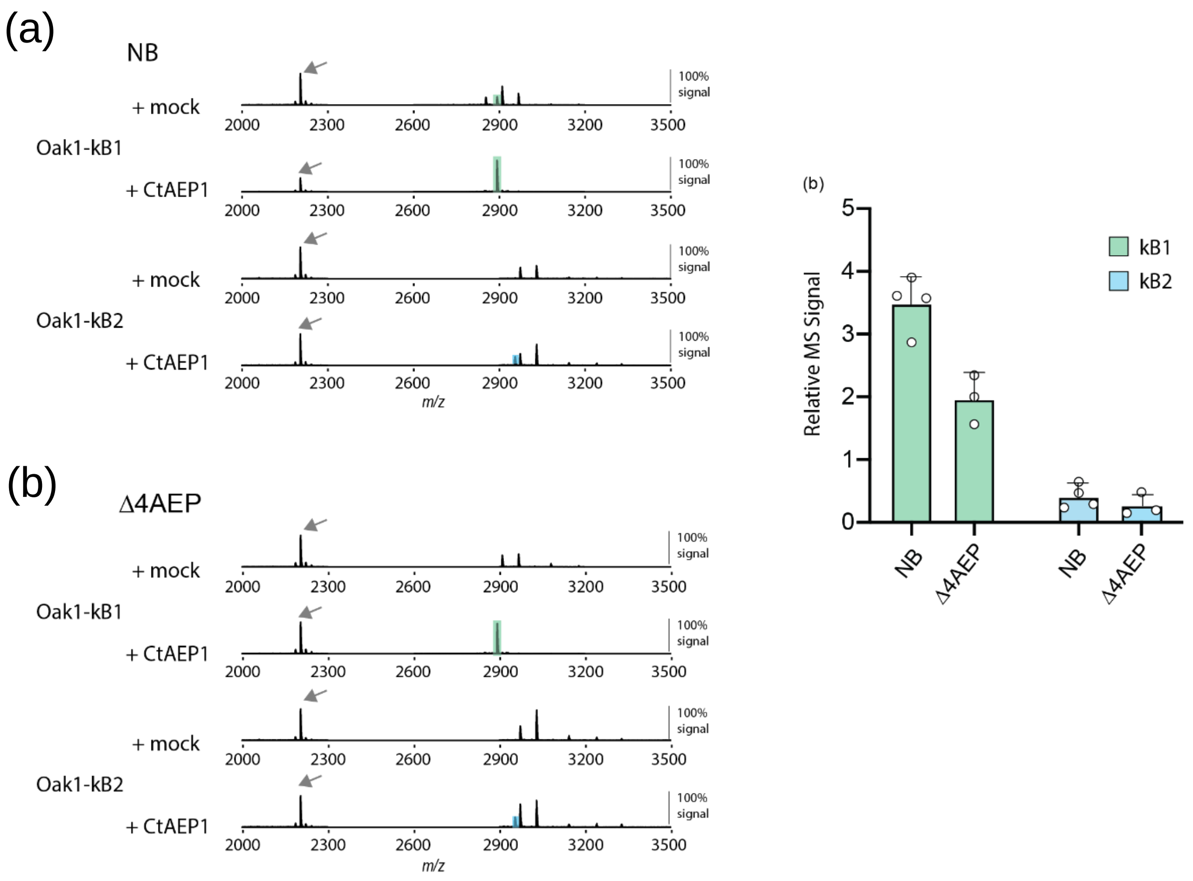


**Figure S2. (a)** Representative MALDI-TOF-MS of kB1 and kB2 accumulated in wild type *N. benthamiana* (NB) and the ΔAEP accession. MS signals for cyclic peptides are highlighted to match the colours in panel b. An arrow indicates the MS signal for the internally spiked peptide control that served to normalise MS signals for relative quantification. **(b)** Mean and SD (n=3 or 4) of relative kB1 and kB2 MS signals detected in crude peptide extracts of infiltrated *N. benthamiana* (NB) and the ΔAEP accession.

(a)

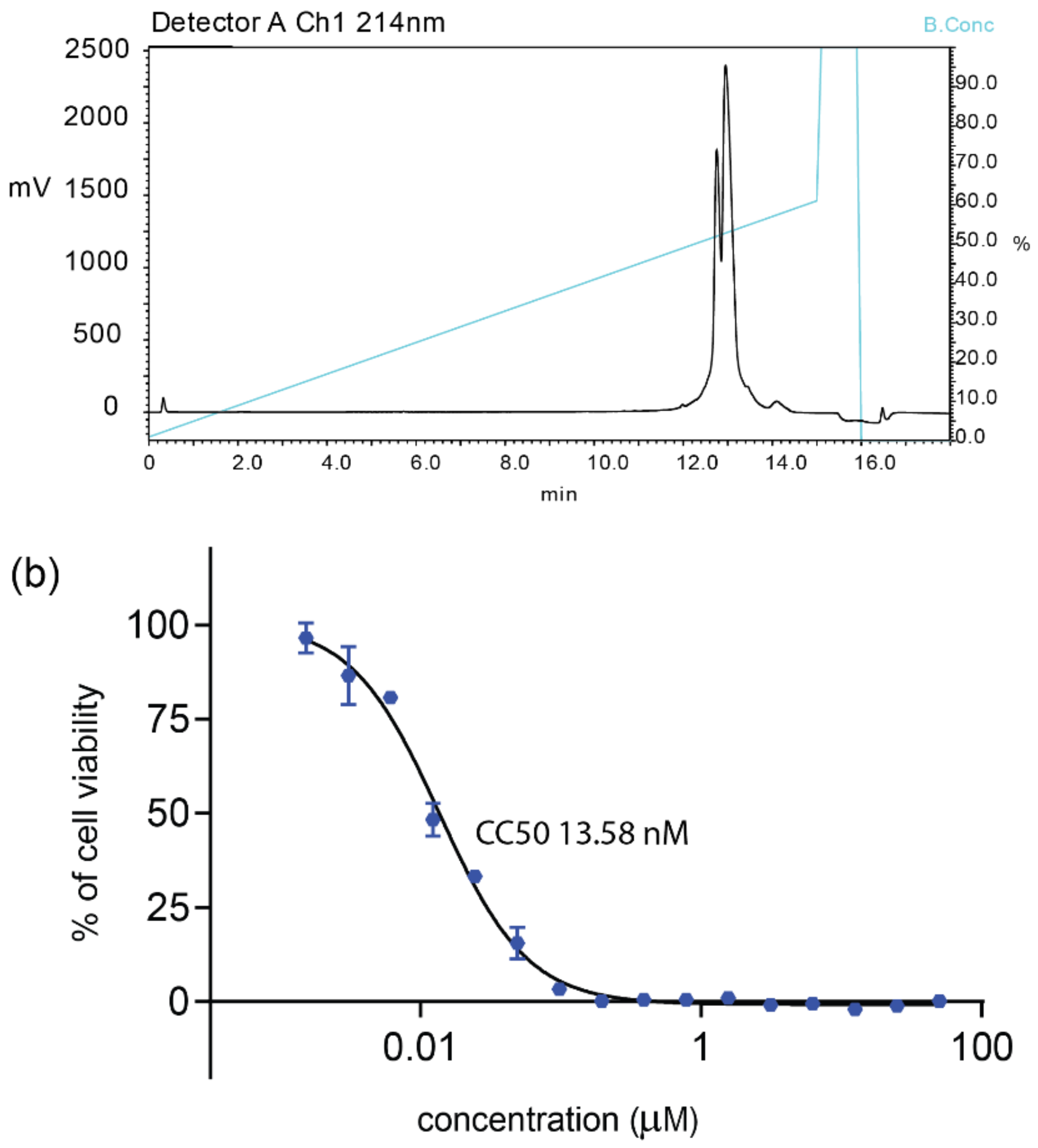


**Figure S3. (a)** HPLC trace of purified Pa1b. The early eluting peak was shown to be Pa1b carrying an oxidised methionine. **(b)** Cytotoxicity of Pa1b against Sf9 cells.

**Oak SFTI**
ATGGCTAAGTTTACCGTGTGCCTTTTGCTCTGCCTTCTCCTCGCTGCTTTTGTTGGAGCTTTCGGATCTGAGCTTTCTGATTCTCACAAGACCACCCTCGTGAACGA
GATCGCTGAGAAGATGCTCCAGAGAAAGATCCTCGATGGTGTTGAGGCTACTCTCGTGACTGATGTGGCAGAGAAGATGTTCCTCAGAAAGATGAAGGCTGAGGCTA
AGACCTCTGAGACTGCTGATCAGGTTTTCCTCAAGCAGCTTCAGCTTAAGGGAAGATGTACCAAGTCTATCCCTCCTATCTGTTTCCCTGATGGACTCCCTTCTCTT
GCTGCTTGA

MAKFTVCLLLCLLLAAFVGAFGSELSDSHKTTLVNEIAEKMLQRKILDGVEATLVTDVAEKMFLRKMKAEAKTSETADQVFLKQLQLK**GRCTKSIPPICFPD**GLPSL
AA*

Primers for SDM PCR to make Oak SFTI_N

SFTI D-N Fwd     GTTTCCCTAATGGACTCCCTTCTCTTGCTGC
SFTI D-N Rev     GTCCATTAGGGAAACAGATAGGAGGGATAGAC

**Oak SFTI KLK4_D**
ATGGCTAAGTTTACCGTGTGCCTTTTGCTCTGCCTTCTCCTCGCTGCTTTTGTTGGAGCTTTCGGATCTGAGCTTTCTGATTCTCACAAGACCACCCTCGTGAACGA
GATCGCTGAGAAGATGCTCCAGAGAAAGATCCTCGATGGTGTTGAGGCTACTCTCGTGACTGATGTGGCAGAGAAGATGTTCCTCAGAAAGATGAAGGCTGAGGCTA
AGACCTCTGAGACTGCTGATCAGGTTTTCCTCAAGCAGCTTCAGCTTAAGGGATTCTGTCAGAGATCTATCCCTCCTATCTGTTTCCCTGATGGACTCCCTTCTCTT
GCTGCTTGA

MAKFTVCLLLCLLLAAFVGAFGSELSDSHKTTLVNEIAEKMLQRKILDGVEATLVTDVAEKMFLRKMKAEAKTSETADQVFLKQLQLK**GFCQRSIPPICFPD**GLPSL
AA*

Primers for SDM PCR to make Oak SFTI KLK4_N

KLK4 D-N Fwd     GTTTCCCTAATGGACTCCCTTCTCTTGCTG
KLK4 D-N Rev     GTCCATTAGGGAAACAGATAGGAGGGATAG

**Oak SFTI KLK5_N**
ATGGCTAAGTTTACCGTGTGCCTTTTGCTCTGCCTTGTCCTCGCTGCTGTTGTTGGAGCTTTCGGATCTGAGCTTTCTGATTCTCACAAGACCACCCTCGTGAACGA
GATCGCTGAGAAGATGCTCCAGAGAAAGATCCTCGATGGTGTTGAGGCTACTCTCGTGACTGATGTGGCAGAGAAGATGTTCCTCAGAAAGATGAAGGCTGAGGCTA
AGACCTCTGAGACTGCTGATCAGGTTTTCCTCAAGCAGCTTCAGCTTAAGGGATATTGTAATAGATCTTATCCTCCTGAATGTTTCCCTAATGGACTCCCTTCTCTT
GCTGCTTGA

MAKFTVCLLLCLVLAAVVGAFGSELSDSHKTTLVNEIAEKMLQRKILDGVEATLVTDVAEKMFLRKMKAEAKTSETADQVFLKQLQLK**GYCNRSYPPECFPN**GLPSL
AA*

**Oak Vc1.1**
ATGGCTAAGTTTACCGTGTGCCTTTTGCTCTGCCTTCTCCTCGCTGCTTTTGTTGGAGCTTTCGGATCTGAGCTTTCTGATTCTCACAAGACCACCCTCGTGAACGA
GATCGCTGAGAAGATGCTCCAGAGAAAGATCCTCGATGGTGTTGAGGCTACTCTCGTGACTGATGTGGCAGAGAAGATGTTCCTCAGAAAGATGAAGGCTGAGGCTA
AGACCTCTGAGACTGCTGATCAGGTTTTCCTCAAGCAGCTTCAGCTTAAGGGATGTTGCTCTGATCCTCGTTGTAATTATGATCATCCTGAAATTTGCTGA

MAKFTVCLLLCLLLAAFVGAFGSELSDSHKTTLVNEIAEKMLQRKILDGVEATLVTDVAEKMFLRKMKAEAKTSETADQVFLKQLQLK**GCCSDPRCNYDHPEIC***

**Oak [N9W]Vc1.1**
ATGGCTAAGTTTACCGTGTGCCTTTTGCTCTGCCTTCTCCTCGCTGCTTTTGTTGGAGCTTTCGGATCTGAGCTTTCTGATTCTCACAAGACCACCCTCGTGAACGA
GATCGCTGAGAAGATGCTCCAGAGAAAGATCCTCGATGGTGTTGAGGCTACTCTCGTGACTGATGTGGCAGAGAAGATGTTCCTCAGAAAGATGAAGGCTGAGGCTA
AGACCTCTGAGACTGCTGATCAGGTTTTCCTCAAGCAGCTTCAGCTTAAGGGATGTTGCTCTGATCCTCGTTGTTGGTATGATCATCCTGAAATTTGCTGA

MAKFTVCLLLCLLLAAFVGAFGSELSDSHKTTLVNEIAEKMLQRKILDGVEATLVTDVAEKMFLRKMKAEAKTSETADQVFLKQLQLK**GCCSDPRCWYDHPEIC***

**Oak_HIIAA**
ATGGCTAAGTTTACCGTGTGCCTTTTGCTCTGCCTTCTCCTCGCTGCTTTTGTTGGAGCTTTCGGATCTGAGCTTTCTGATTCTCACAAGACCACCCTCGTGAACGA
GATCGCTGAGAAGATGCTCCAGAGAAAGATCCTCGATGGTGTTGAGGCTACTCTCGTGACTGATGTGGCAGAGAAGATGTTCCTCAGAAAGATGAAGGCTGAGGCTA
AGACCTCTGAGACTGCTGATCAGGTTTTCCTCAAGCAGCTTCAGCTTAAGGGACTCCCTGTTTGCGGAGAGACTTGTGTTGGAGGAACTTGCAACACTCCTGGATGC
ACTTGTTCTTGGCCTGTGTGTACTAGAAACCATATTATCGCTGCTTGA

MAKFTVCLLLCLLLAAFVGAFGSELSDSHKTTLVNEIAEKMLQRKILDGVEATLVTDVAEKMFLRKMKAEAKTSETADQVFLKQLQLK**GLPVCGETCVGGTCNTPGC
TCSWPVCTRN**HIIAA*
**PA1**

ATGGCTTCTGTTAAGCTTGCTTCTCTGATCGTGCTGTTCGCTACCCTTGGTATGTTCCTGACTAAGAACGTGGGTGCTGCTTCTTGCAATGGTGTGTGCTCTCCTTT
CGAAATGCCTCCTTGTGGTACTAGCGCTTGCAGATGCATTCCTGTGGGTCTTGTTGTGGGATACTGCAGAAATCCTAGCGGTGTGTTCCTGAGGACTAACGATGAGC
ATCCTAACCTGTGCGAGTCCGATGCTGATTGCAGAAAGAAGGGTTCCGGTAACTTCTGCGGTCACTACCCTAACCCTGATATCGAGTACGGTTGGTGCTTCGCTTCT
AAGTCTGAGGCTGAGGATTTTTTCAGCAAGATTACCCCTAAGGATCTGCTGAAGTCCGTGTCTACTGCTTAG

MASVKLASLIVLFATLGMFLTKNVGA**ASCNGVCSPFEMPPCGTSACRCIPVGLVVGYCRNPSG**VFLRTNDEHPNLCESDADCRKKGSGNFCGHYPNPDIEYGWCFAS
KSEAEDFFSKITPKDLLKSVSTA*

**Figure S4.** Gene sequences ordered as dsDNA gene blocks and primers designed for site directed mutagenesis experiments.

**Supplementary Method 1**

**pGEMT-NbAEP construction**

| Product name | For-Name | Forward-Seq | Rev-Name | Rev-Seq | Size (bp) |
|---|---|---|---|---|---|
| pGEMT-MOD | pGEMT-AarI_F | TGCAGTTTTTTTGCAGGTGGGCGAATCAC TAGTGCGGCCGCCTGCAGG | pGEMT-AarI_R | GTTCAATTTTTGCAGGTGGCGCTCAT CCCGCGGCCATGGCGGC | 3003 |
| Nb-PROD1 | pTG_5ter mAarI_F | CACCTGCAAAAATTGAACAAAGCACCAG TGGTCTAG | NbAEP1p TGv2_R | ACCGATCCGTACCTCTATGCTGCAC CAGCCGGGAATCG | 112 |
| Nb-PROD2 | NbAEP1p TGv2_F | GCATAGAGGTACGGATCGGTGTTTTAGA GCTAGAAATAGC | NbAEP2p TGv2_R | GGCAGATGTTTGTCACGCATTGCAC CAGCCGGGAATCG | 193 |
| Nb-PROD3 | NbAEP2p TGv2_F | ATGCGTGACAAACATCTGCCGTTTTAGA GCTAGAAATAGC | NbAEP3p TGv2_R | CAGCTGCTCTCTACAGCGTTTGCACC AGCCGGGAATCG | 193 |
| Nb-PROD4 | NbAEP3p TGv2_F | AACGCTGTAGAGAGCAGCTGGTTTTAGA GCTAGAAATAGC | NbAEP4p TGv2_R | GGCCTTGGATTCAGCTCACTTGCAC CAGCCGGGAATCG | 193 |
| Nb-PROD5 | NbAEP4p TGv2_F | AGTGAGCTGAATCCAAGGCCGTTTTAGA GCTAGAAATAGC | pTG_3ter mAarI_R | CACCTGCAAAAAAACTGCACCAGCC GGGAATC | 188 |

>pTG_mid

GTTTTAGAGCTAGAAATAGCAAGTTAAAATAAGGCTAGTCCGTTATCAACTTGAAAAAGTGGCACCGAGTCGGTGCAACAAAGC ACCAGTGGTCTAGTGGTAGAATAGTACCCTGCCACGGTACAGACCCGGGTTCGATTCCCGGCTGGTGCA

Method:

1. Products were amplified using Phusion polymerase (ThermoFisher Scientific) as per manufacturer’s protocol. Annealing temperature for first five cycles was 55°C for 10s followed by thirty cycles as a two-step cycle of 98°C for 10s and 72°C for the appropriate recommended extension time. Addition of 3% DMSO final concentration to pGEMT-MOD reaction was beneficial. Template for pGEMT-MOD was linearized pGEM-T Easy (Promega), and template for all other reactions was the pTG-mid dsDNA fragment synthesized by Integrated DNA Technologies.

2. All products were run on either a 1% (pGEMT-MOD) or 2% (all others) agarose TBE gel, visualized with SybrSafe stain (Invitrogen) and cut out for purification (Macherey-Nagel NucleoSpin Gel and PCR Purification kit).
3. All products were quantified using spectrophotometry, and calculated amounts equal to 0.02pmol each were added to 2x NEBuilder HiFi (NEB) and assembled as per manufacturer's protocol.
4. The resulting plasmid yields an AarI digestion fragment equal to 777bp in size with the structure:

*ATTG overhang*+tRNAgly:**NbAEP1-gRNA**:tRNA-gly:**NbAEP2-gRNA**:tRNA-gly:**NbAEP3-gRNA**:tRNA-gly:**NbAEP4-gRNA**:tRNA-gly+*GTTT overhang*

**Supplementary Method 2**

**Genotyping ΔAEP plants**

| Product name | Expected size (bp) | Name | Sequence |
|---|---|---|---|
| NbAEP1 | 534 | NbAEP1_probeF | TAGATATCACTGTCGTATATGGAGG |
| | | NbAEP1_probeR | ATAGGTTCCCCAGCTACTCTC |
| NbAEP2 | 604 | NbAEP2_probeF | AGTAGCGGTTTATGAGCATG |
| | | NbAEP2_probeR | GCTGTAACATGATGCCCTG |
| NbAEP3 | 311 | NbAEP3_probeF | AAGCATGCATCTGGGACATA |
| | | NbAEP3_probeR | AACAGCATACAAGTCACCCA |
| NbAEP4 | 381 | NbAEP4_probeF | CTGGGTTACTGATCGATGCT |
| | | NbAEP4_probeR | GAGGCACCATTATTGATGCC |

**Method for CAPS:**

1. gDNA purified from young leaves using Jena Science Plant DNA kit.
2. Reaction mixes were set up with gDNA a sample from two isolated plants grown many generations after initial selection. Amplicons amplified with DNA Taq Polymerase (Invitrogen) per manufacturer's protocol.
3. 10µL of PCR product was assembled into restriction digest reactions as per NEB Restriction Digest protocol (NEB) at 30µL volume. Control (undigested) product was also assembled into buffer+dilutent mock reactions minus enzyme and incubated alongside enzyme-containing reactions. The enzymes used were: NbAEP1 (AlwI), NbAEP2 (PspGI), NbAEP3 (PvuII), NbAEP4 (StyI).
4. Products were run on a gel to visualize CAPS markers, with cleaved products representing wild-type alleles and full-length probes representing alleles where the restriction enzyme site was obliterated.

**Method for Sanger sequencing:**

1. gDNA purified from young leaves using Jena Science Plant DNA kit.
2. Reaction mixes were set up with gDNA. Amplicons amplified with Phusion polymerase (ThermoFisher Scientific).
3. Products were run on a gel to visualize bands, and for gel purification for Sanger sequencing. Product sizes are given in the primers table.
4. Primers used for sequencing were the forward primers for each NbAEP probe.

**Table S1. Genes significantly upregulated and downregulated greater than 2-fold.**

| GeneID | logFC | logCPM | F | PValue | Blast-Hit-Accession | Human-Readable-Description |
|---|---|---|---|---|---|---|
| Niben101Scf02659g01005.1 | -3.430585879 | 3.622925739 | 90.71901253 | 0.000195837 | sp\|A8H1G3\|G6PI_SHEPA | Glucose-6-phosphate isomerase |
| Niben101Scf04539g04014.1 | -3.28908874 | 3.947356753 | 121.6809836 | 9.55659E-05 | sp\|P49043\|VPE_CITSI | Vacuolar-processing enzyme |
| Niben101Scf04675g08014.1 | -2.946428049 | 5.849135063 | 64.58170801 | 0.000443867 | sp\|P49043\|VPE_CITSI | Vacuolar-processing enzyme |
| Niben101Scf18356g00003.1 | -2.913219846 | 4.700707579 | 50.47443889 | 0.000795079 | sp\|P49043\|VPE_CITSI | Vacuolar-processing enzyme |
| Niben101Scf07493g04001.1 | 2.043953945 | 4.082535494 | 63.9637378 | 0.000454159 | AT5G42830.1 | HXXXD-type acyl-transferase family protein LENGTH=450 |
| Niben101Scf04273g00001.1 | 2.088021671 | 4.87781374 | 7.48930441 | 0.040191691 | sp\|Q11PV4\|METE_CYTH3 | 5-methyltetrahydropteroyltriglutamate--homocysteine methyltransferase |
| Niben101Scf03147g09006.1 | 2.288482969 | 4.599855983 | 8.415615305 | 0.03305199 | sp\|Q9SR36\|GSTU8_ARATH | Glutathione S-transferase U8 |
| Niben101Scf01789g04010.1 | 2.436064447 | 5.133006085 | 7.813826473 | 0.037463521 | sp\|Q15782\|CH3L2_HUMAN | Chitinase-3-like protein 2 |
| Niben101Scf01481g01006.1 | 2.648302619 | 4.772189075 | 28.28017701 | 0.002981915 | sp\|Q9FLS0\|FB253_ARATH | F-box protein |
| Niben101Scf01970g01015.1 | 2.6541105 | 3.637694869 | 28.41805886 | 0.002950029 | sp\|Q0VBY0\|CMC4_BOVIN | Cx9C motif-containing protein 4 |
| Niben101Scf07242g01001.1 | 2.791907417 | 5.461647938 | 7.293642233 | 0.04197315 | AT3G28740.1 | Cytochrome P450 superfamily protein LENGTH=509 |
| Niben101Scf01660g00007.1 | 2.981822846 | 4.909884544 | 7.812794329 | 0.037471779 | AT4G14090.1 | UDP-Glycosyltransferase superfamily protein LENGTH=456 |
| Niben101Scf02217g07009.1 | 3.109945068 | 6.206317215 | 8.649909024 | 0.031530882 | sp\|Q4FRX3\|MAO1_PSYA2 | NAD-dependent malic enzyme |
| Niben101Scf02349g03001.1 | 3.394246764 | 7.402016387 | 20.95584041 | 0.005703932 | sp\|A7NY33\|PER4_VITVI | Peroxidase 4 |
| Niben101Scf01237g09002.1 | 3.790827628 | 3.3046639 | 51.71549607 | 0.000751034 | sp\|Q9FFB8\|CHX3_ARATH | Cation/H(+) antiporter 3 |
| Niben101Scf03886g04003.1 | 3.868166949 | 4.323765449 | 20.76272895 | 0.005816618 | sp\|Q9FED2\|HA22E_ARATH | HVA22-like protein e |
| Niben101Scf14427g00009.1 | 5.106760145 | 4.85019011 | 19.49419786 | 0.00664011 | 0 | Unknown protein |
